# Quantifying variation in bacterial reproductive fitness with and without antimicrobial resistance: a high-throughput method

**DOI:** 10.1101/2020.05.13.093807

**Authors:** Pascal M. Frey, Julian Baer, Judith Bergadà Pijuan, Conor Lawless, Philipp K. Bühler, Roger D. Kouyos, Katherine P. Lemon, Annelies S. Zinkernagel, Silvio D. Brugger

## Abstract

**Background:** To evaluate changes in reproductive fitness of bacteria, e.g., after acquisition of antimicrobial resistance, a low-cost high-throughput method to analyse bacterial growth on agar is desirable for broad usability, including in low-resource settings.

**Method:** In our bacterial quantitative fitness analysis (baQFA), cultures are spotted in a predefined array on agar plates and photographed sequentially while growing. These time-lapse images are analysed using a purpose-built open source software to derive normalised image intensity values for each culture spot. Subsequently, a Gompertz growth model is fitted to these optical intensity values of each culture spot and fitness is calculated from parameters of the model. For image segmentation validation, we investigated the association between normalised intensity values and colony-forming unit (CFU) counts. To represent a range of clinically important pathogenic bacteria, we used different strains of *Enterococcus faecium, Escherichia coli* and *Staphylococcus aureus*, with and without antimicrobial resistance. Relative competitive fitness (RCF) was defined as the mean fitness ratio of two strains growing competitively on one plate.

**Results:** baQFA permitted the accurate construction of growth curves from bacteria grown on semisolid agar plates and fitting of Gompertz models: Normalised image intensity values showed a strong association with the total CFU/ml count per spotted culture (p<0.001) for all strains of the three species. Bacterial QFA showed relevant reproductive fitness differences between individual strains, suggesting substantial higher fitness of methicillin-resistant *S. aureus* JE2 than Cowan (RCF 1.60, p<0.001). Similarly, the vancomycin-resistant *E. faecium* ST172b showed higher competitive fitness than susceptible *E. faecium* ST172 (RCF 1.72, p<0.001).

**Conclusion:** Our baQFA adaptation allows detection of fitness differences between our bacterial strains, and is likely to be applicable to other bacteria. In the future, baQFA may help to estimate epidemiological antimicrobial persistence or contribute to the prediction of clinical outcomes in severe infections at a low cost.

## INTRODUCTION

Reproductive fitness is the ultimate target of evolution, and in general, no cells can afford a reduction in fitness. Therefore, quantifying changes in fitness is highly informative about the evolutionary potential of cells.

Antimicrobial-resistant strains of pathogenic bacteria with the potential to cause severe infections in humans are widely regarded as a threat to public health. Although research is often focused on the development and acquisition of antimicrobial resistance, it is generally thought that most acquired antimicrobial resistances come with a cost in reproductive fitness, and thus they are expected to disappear if selection pressure from antibiotics is reduced [1]. However, epidemiologic evidence to support this assumption is scarce, and the routine use of methods to screen for changes in the reproductive fitness of emerging antimicrobial resistant bacterial strains are not widely applied, possibly due to a lack of availability of high-throughput and low-cost methods, especially in low-resource settings that show a high prevalence of antibiotic resistance [2].

Furthermore, a high-throughput method to accurately determine reproductive fitness in bacteria despite an expectedly high in-strain phenotypic variability, may not only be an important instrument in predicting the epidemiologic persistence of resistant strains [1], but might also provide a useful insight regarding clinical outcomes of severe infections in general. In order to investigate the fitness of bacteria in smaller laboratories and in settings with limited resources, a low-cost high-throughput method is needed to cope with the large amount of data required for routine analysis.

A similar requirement has previously driven the design of automated synthetic genetic array (SGA) methods, where interactions of genes, usually in yeast as a model eukaryotic cell, are studied in large scale while allowing interaction between mutants while growing [3]. Among other approaches, this need has also led to the development of an automated high-throughput quantitative fitness analysis (QFA) method for yeast, which was used as a eukaryotic model for the investigation of reproductive fitness of human cell mutations associated with telomere capping [4]. In QFA, an array of yeast cultures (e.g., 8 by 12, or 16 by 24) with different mutants is spotted onto a rectangular agar plate in a predefined pattern, where every single culture spot is then growing in competition to its neighbour mutant. During growth, time-lapse photographs of the agar plate with its cultures are taken in a defined time interval. These images are then processed by a purpose-built open source software, which derives intensity values for each culture spot [5], as a surrogate measure for each spot’s microbial population density. These intensity values are subsequently analysed using a designated R package [6], which fits a logistic growth model over measured time-lapse values and derives fitness from parametrization of the mathematical model [4,5].

A similar method for quantifying the fitness of *Escherichia coli* mutants, “Colony-live”, has also been developed [7]. This method is well validated and accurate; however, it relies on one dedicated plate scanner per plate, with a scanning illumination that is placed to measure absorbance of growing colonies in a way that is unsuitable for measuring the reflection of light from the colonies as required with regular photo cameras. The dependence of the Colony-live method on expensive specialized equipment leaves a need for a low-cost and flexible method to be automated with consumer electronics.

However, the principles of the QFA method are considered to be applicable to bacteria as well [4]. The QFA setup is flexible and the time-lapse photo analysis algorithm is designed to read normal photographs from a consumer electronics DSLR camera. Thus, we aimed to adapt the QFA method for use with *Staphylococcus aureus* and other important bacterial pathogens in human medicine, using low-cost consumer products for basic QFA automation and data acquisition.

We hypothesized that an adaptation of the QFA method to bacteria would yield valid bacterial reproductive fitness results accurate enough to detect differences between strains with different antimicrobial resistance properties despite an expectedly high random biological variation.

## METHODS

### Quantitative fitness analysis (QFA) and bacterial pathogens

Our bacterial QFA (baQFA) adaptation consisted of cultures of the microbe of interest to be spotted onto solid agar medium in a rectangular array of small neighbouring individual culture inoculations. The growth of these neighbouring inocula was then followed over time by automated high-resolution time-lapse photography. The images were processed using our custom-built open source software “BaColonyzer” to derive normalised intensity values. These were subsequently fitted to a mathematical growth model for an accurate estimation of growth parameters, from which reproductive fitness was derived. In our adaptation of the method for bacteria, we used a Gompertz growth model for derivation of reproductive fitness parameters (**Figure 1**).

**Figure 1:**
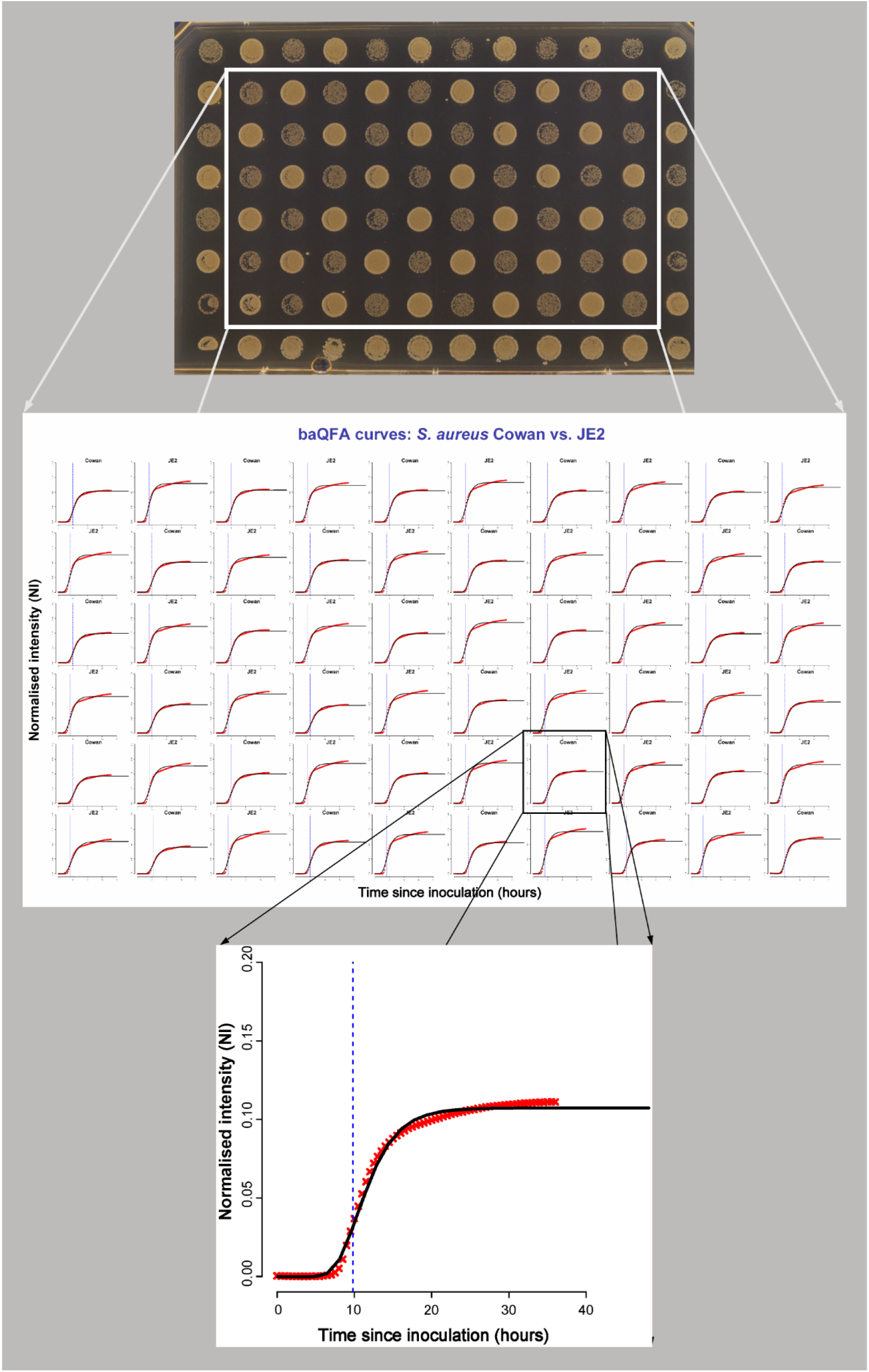
Construction of growth curves and fitness measures from time-lapse photos of *S. aureus* strains Cowan and JE2

To represent a range of important bacterial pathogens in humans, we decided to assess the accuracy of the baQFA method adaptation using mainly *Staphylococcus aureus* (methicillin resistant JE2; methicillin-susceptible Cowan), but also *Enterococcus faecium* (ST172, ST172b), and *Escherichia coli* DH5α (**Table 1**).

**Table 1:**
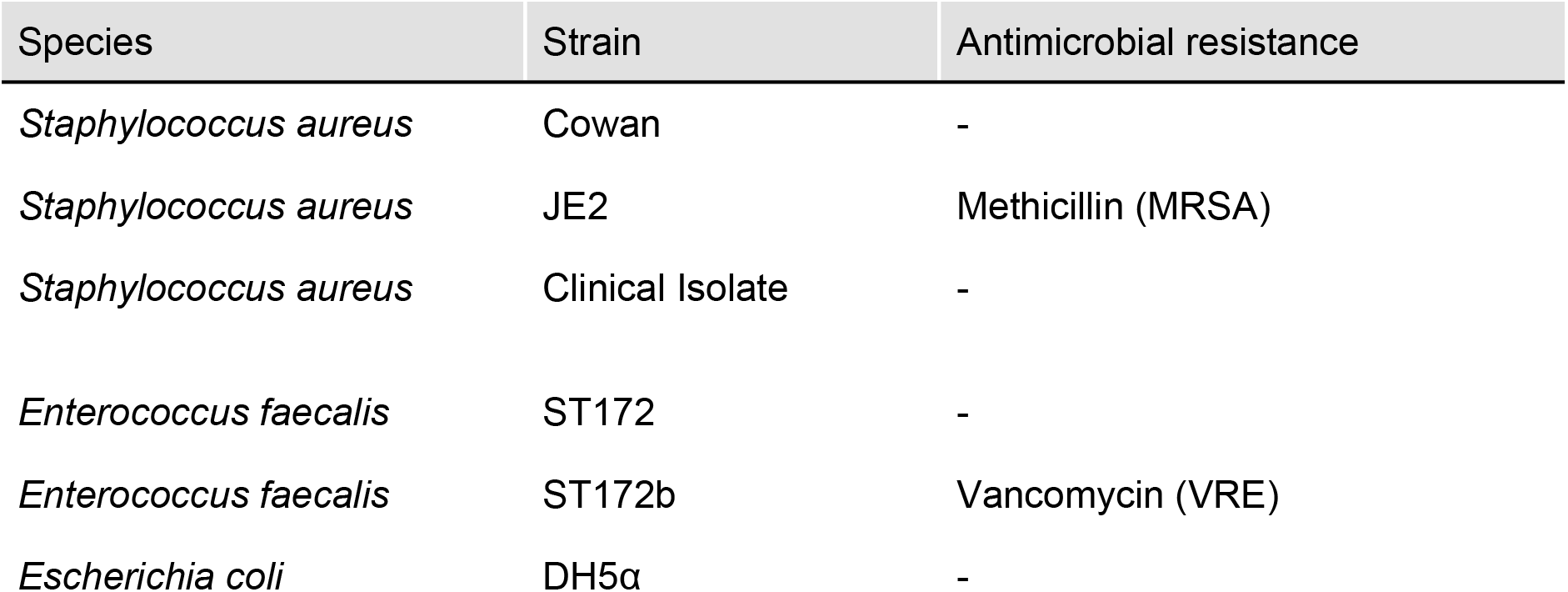
overview of bacterial strains used for baQFA with their antimicrobial resistance.

| Species | Strain | Antimicrobial resistance |
| --- | --- | --- |
| *Staphylococcus aureus* | Cowan | - |
| *Staphylococcus aureus* | JE2 | Methicillin (MRSA) |
| *Staphylococcus aureus* | Clinical Isolate | - |
| *Enterococcus faecalis* | ST172 | - |
| *Enterococcus faecalis* | ST172b | Vancomycin (VRE) |
| *Escherichia coli* | DH5α | - |

### Microbial culture array and growth conditions

In preparation for baQFA, the bacterial strains of interest were streaked out on Columbia sheep blood agar (CSBA) plates and incubated at 37°C for 18 to 24 h. Freshly grown strains were then harvested from the plate with sterile loops and resuspended in Phosphate Buffered Saline (PBS). After vortexing, the solution was diluted to an optical density (OD_600nm_) of 0.1 (±0.01).

To prepare the final dilution, 1ml of this 0.1 OD_600_ solution was diluted in 40ml PBS (or for lower quantities 0.5 ml in 20 ml to receive the same final concentration). We then added 200 µl of this final fresh bacterial suspension to each well of a 96-well plate in the desired pattern (e.g., a checkerboard grid: placing each tested strain in direct neighbourhood of the competing second strain tested for a 2-strain array).

Using a 96-pin replica plater, bacteria were finally transferred from the 96-well plate to a rectangular single-well BHI agar plate (20 ml agar medium, ThermoScientific OmniTray Single Well w/ lid). The BHI agar plate was then put into the baQFA imaging setup and incubated at 37°C ambient air (no additional CO_2_). No antibiotics were added to any of the plates.

### Culture image capturing

Image capturing was automated using a LEGO® Mindstorms® EV3 robot, assembled into a purpose-built PVC box within a standard incubator. For plate illumination, we used a consumer electronic cold white LED stripe.

Mounted on a Manfrotto® table top tripod (Kit 209+492), we used a Canon® EOS D650 DSLR camera with a Canon® EF 40mm f/2.8 STM lens for imaging. Images were taken at a resolution of 5184×3456 pixels, with ISO 400, aperture f/22 and exposure of 2.5 seconds in manual mode. The camera shutter was controlled through the LEGO® Mindstorms® robot and programmed to release every 10, 20 or 30 minutes after the robot had opened the plate lid (to reduce reflections and condensation artefacts; lid is needed to keep agar from drying).

For further automated processing, each captured image file was renamed to reflect the exact time and date of capturing using Canon® image metadata.

### Image segmentation and normalised intensity values of bacterial growth

For the derivation of normalised intensity (NI) values of each culture spot on the time lapse image series, the open-source software BaColonyzer was created using Python3. Inspired by Colonyzer [5], the new algorithm is much faster and adapted to be more robust to handle bacterial colonies. BaColonyzer imports a grayscale version of the images using OpenCV (OpenCV, 2015. Open Source Computer Vision Library). The NI is based on the intensity values of the pixels. BaColonyzer finds the location of the plate by creating an artificial template of the grid, which is then matched to the last image in the time-lapse series. This allows removal of the borders and other pixels that might introduce noise or artefacts. Otsu’s binarization is used to identify the culture spot areas and the agar areas. For a proper NI quantification, BaColonyzer divides the agar plate into smaller patches that contain one colony spot. Next, it normalises the intensity values of each patch by subtracting the mean intensity of the agar in this patch, thus correcting for colour differences within and between images. NI of each colony is finally computed as the sum of all intensity values of the patch, divided by the number of pixels. Furthermore, BaColonyzer includes an optional normalization that accounts for the camera settings, improving the comparison of various sets of images. A visualisation of the conceptual design has been given in **Figure 2**, while the algorithm is described in more detail in **Supplement A**. BaColonyzer is available as a package from PyPI for Python3, which ensures a fast and easy installation. Documentation, instructions and the source code can be found online on GitHub (https://github.com/baQFA).

**Figure 2:**
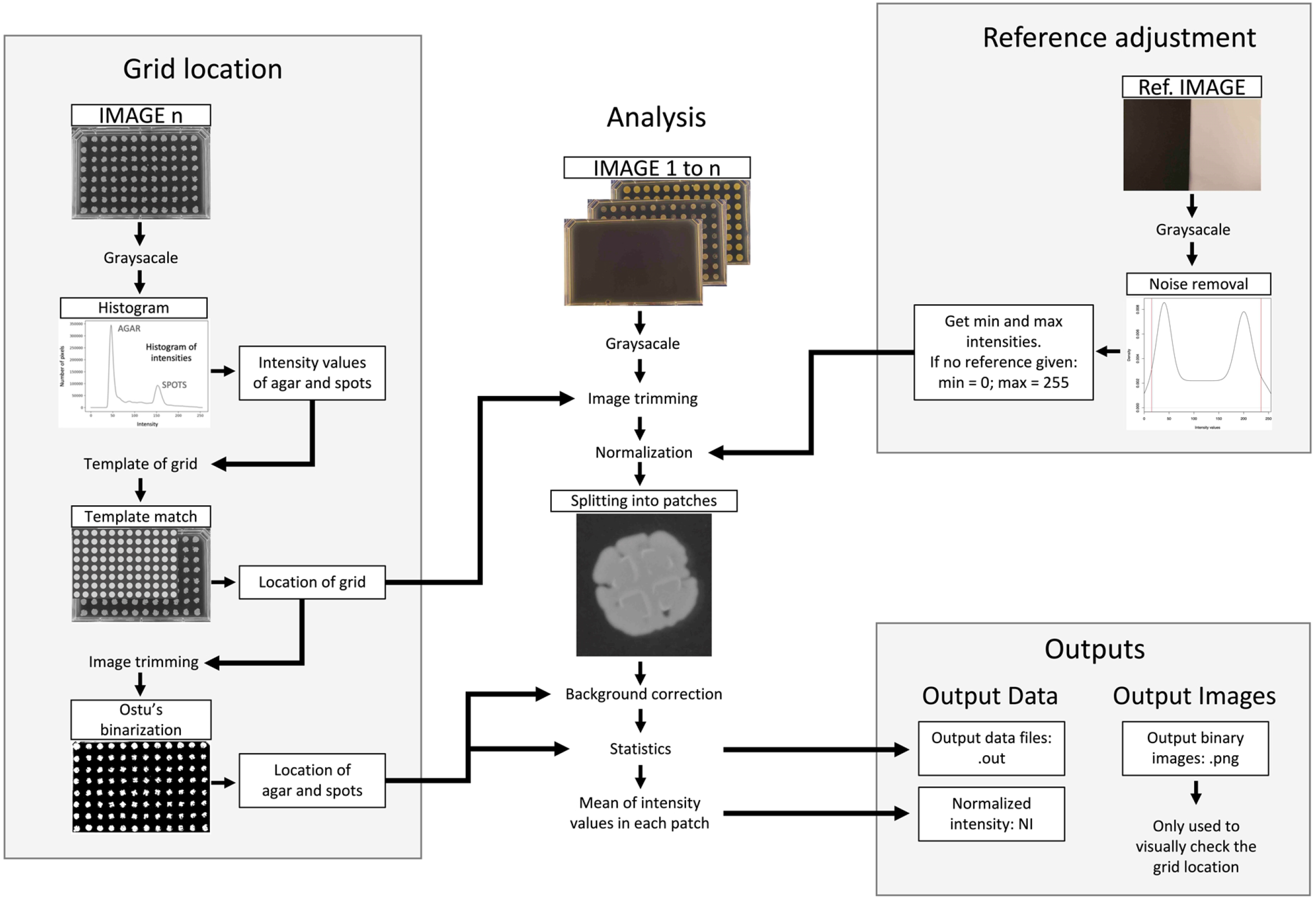
Conceptual design of the BaColonyzer algorithm

To investigate the assumption of our derived intensity values as a surrogate for the true bacterial population density in a culture spot, we examined the association of computationally derived NI measures and bacterial colony-forming unit (CFU) counts of the same culture spot. For this, an agar biopsy of the culture spot was vortexed in PBS to suspend the bacteria on it in solution. From this suspension we performed a dilution series on CSBA plates to estimate the total number of bacterial cells (per ml) on a culture spot. These CFU counts were then matched to the BaColonyzer NI value of the same spot taken before biopsy.

After logarithmic transformation of both the NI values as well as the CFU counts, we used linear regression to determine the association between NI values and CFU counts.

To compare the performance of the original Colonyzer software for yeast to our newly built BaColonyzer for image segmentation of bacterial growth, we ran the same culture spot images through each software and compared intensity values for the same spots with the Bland-Altman method [8].

### Growth model parametrization and fitness derivation

In order to estimate fitness, we used a variation of the Gompertz growth model [9], which was fitted to the normalised intensity values over all time points for each culture spot. Fitness measures were then derived from parametrization of the model. Our variation of the Gompertz model can be defined by a carrying capacity parameter K, absolute maximum growth rate MGR and the time t to MGR (T_max_), where the population size is W(t) [9]:

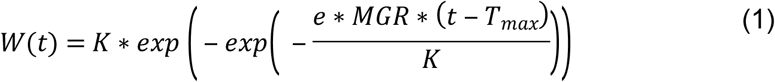

Because of the asymmetrical shape of the bacterial growth curves, the fitting of a logistic model - like in the original QFA method with yeast - is imperfect. We decided to switch to a Gompertz model, which is well established for bacterial growth [7, 9].

Following Addinall et al. [4], we calculated fitness from two measures of growth: a more simplistic non-logarithmic maximum absolute growth rate (MGR), and the time to the maximum absolute growth rate (T_max_), where fitness was defined as MGR divided by T_max_ (Formula 2). With these adaptations, a strain with a larger growth rate and a shorter time to maximum growth would reflect a higher reproductive fitness. Similar fitness estimates have been established previously [1,4,7,10].

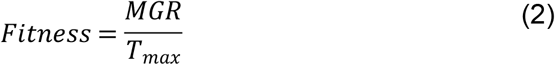

The original QFA method included an R package (“qfaR”) for model fitting and calculation of fitness measures. This R package has been renamed “BaQFA” and adapted to fit the need of using it with bacteria, including the new Gompertz model and fitness measure described above. The BaColonyzer, BaQFA and the LEGO®◻ Mindstorms®◻ program are open-source and available on GitHub (https://github.com/baQFA).

### Fitness comparisons and statistical analyses of fitness

Derived fitness measures from BaQFA were aggregated as mean (SD) fitness, and a density function to represent the fitness distribution per tested strain was visualised using extended violin plots. Since the application of fitness is relative [1], we introduced a relative competitive fitness measure to better quantify differences between two strains when grown in a grid pattern of direct competition to each other on one plate. For this, the ratio of mean fitness was used, with Fieller’s method to calculate exact 95% confidence intervals [11]. Results were considered statistically significant at an alpha level of 0.05. All fitness analyses were performed using R 3.6 (R Core Team. R Foundation for Statistical Computing; 2019).

## RESULTS

Overall, we have analysed the fitness from 719 single culture spots for this adaptation of the bacterial quantitative fitness analysis (baQFA) method. After the adjustments to the culture spotting routine and the computational processing using the new BaColonyzer software as well as BaQFA with Gompertz model fitting, we were able to apply the modified baQFA method to all *S. aureus* and *E. faecium* strains. For *E. coli* we successfully derived intensity values and their correlation to CFU counts, ensuring full functionality with these Gram-negative bacteria.

### Derived normalised intensity values and colony-forming unit (CFU) counts

We found a strong association between CFU counts from single culture spots and the respective normalised intensity measurement from BaColonyzer taken just before determination of CFU counts. Associations for all examined strains were approximately log-linear (**Figure 2**). Bland-Altman plots comparing measurements of the same bacterial cultures with Colonyzer software for yeast and the new BaColonyzer suggested that BaColonyzer measures the normalised intensity on a larger range, with smaller values in low intensity cultures and larger values in high intensity cultures (**Supplement B**).

### Reproductive fitness with and without antibiotic resistance

#### *S. aureus* strains *Cowan* and JE2

As proof of principle that we could detect fitness differences of strains with varying antibiotic resistance properties, we compared two strains of *S. aureus*. Different fitness was observed for the tested strains of *S. aureus*, with JE2 showing higher fitness (**Table 2**), and a broader biologic variability of fitness than Cowan (**Figure 3**).

**Table 2:** Absolute and relative competitive fitness when growing in competition.

| Species | Strain | Mean fitness* (SD) | RCF (95%-CI) | p-value |
| --- | --- | --- | --- | --- |
| *Staphylococcus aureus* | Cowan | 1.3522 (0.2292) | reference |  |
| *Staphylococcus aureus* | JE2 | 2.1725 (0.3705) | 1.61 (1.53 - 1.69) | < 0.0001 |
| *Enterococcus faecalis* | ST172 | 0.2605 (0.0607) | reference |  |
| *Enterococcus faecalis* | ST172b | 0.4473 (0.2153) | 1.72 (1.53 - 1.91) | < 0.0001 |

\*given as mNI/h<sup>2</sup>
Abbreviations: SD = standard deviation; RCF = relative competitive fitness; CI = confidence interval; NI = normalised intensity

**Figure 3:**
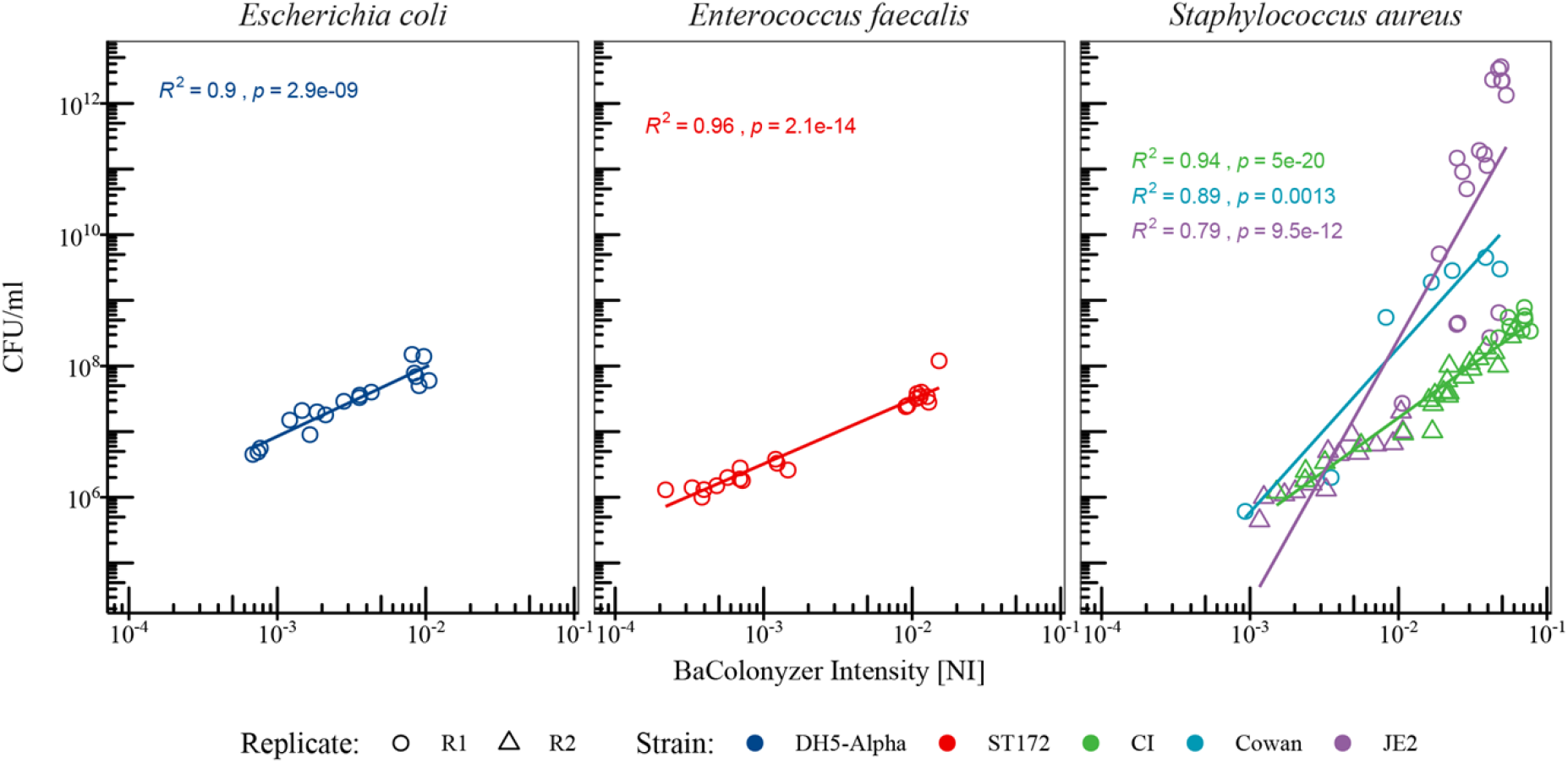
Correlation of colony forming units (CFU) and BaColonyzer derived intensity measures

When grown competitively in the absence of antibiotics, we found strong evidence for a lower fitness of the methicillin-susceptible Cowan strain compared to the methicillin-resistant JE2 strain (overall relative competitive fitness [RCF] 1.61, CI 1.53 to 1.69, p < 0.001). Between three replications of baQFA the RCF varied from 1.50 (CI 1.38 to 1.63) to 1.58 (CI 1.46 to 1.72) to 1.73 (CI 1.64 to 1.84) for replicate 1, 2 and 3, respectively. The presence of the Cowan strain on a grid layout increased fitness for JE2, while JE2 decreased the fitness of Cowan, a sign of possible interaction (**Figure 3**).

#### *E. faecium* strains ST172 and ST172b

When grown in competition we observed a higher fitness of the vancomycin-resistant *E. faecium* strain ST172b compared to the genetically very similar vancomycin-susceptible strain ST172 (**Figure 4**). The ST172b was on average 1.72 times more fit than ST172 (**Table 2**).

**Figure 4:**
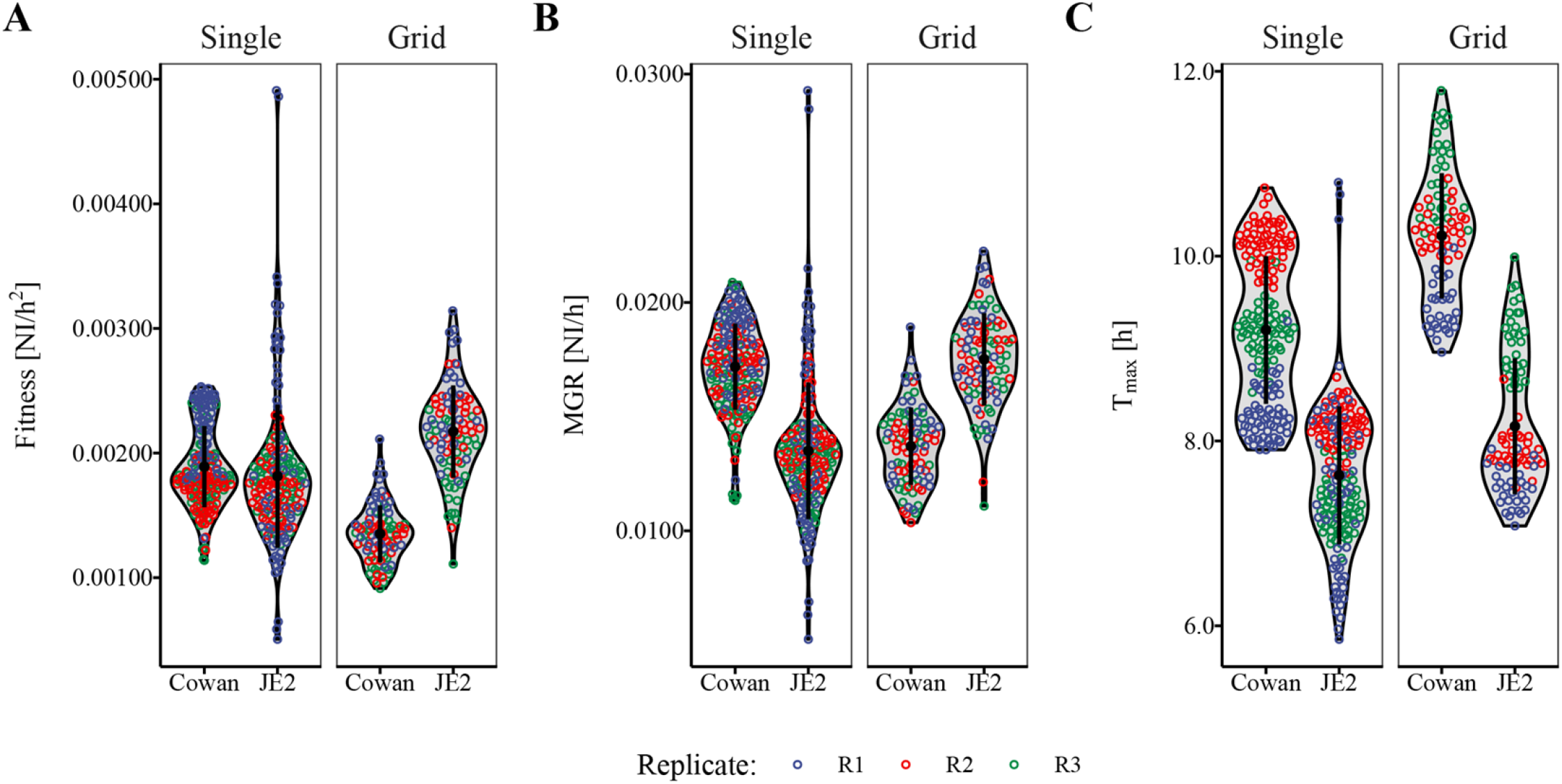
Fitness of methicillin-resistant *S. aureus* (MRSA) strain JE2 compared to the methicillin-susceptible strain Cowan when grown on agar either alone or together in a grid pattern.

**Figure 5:**
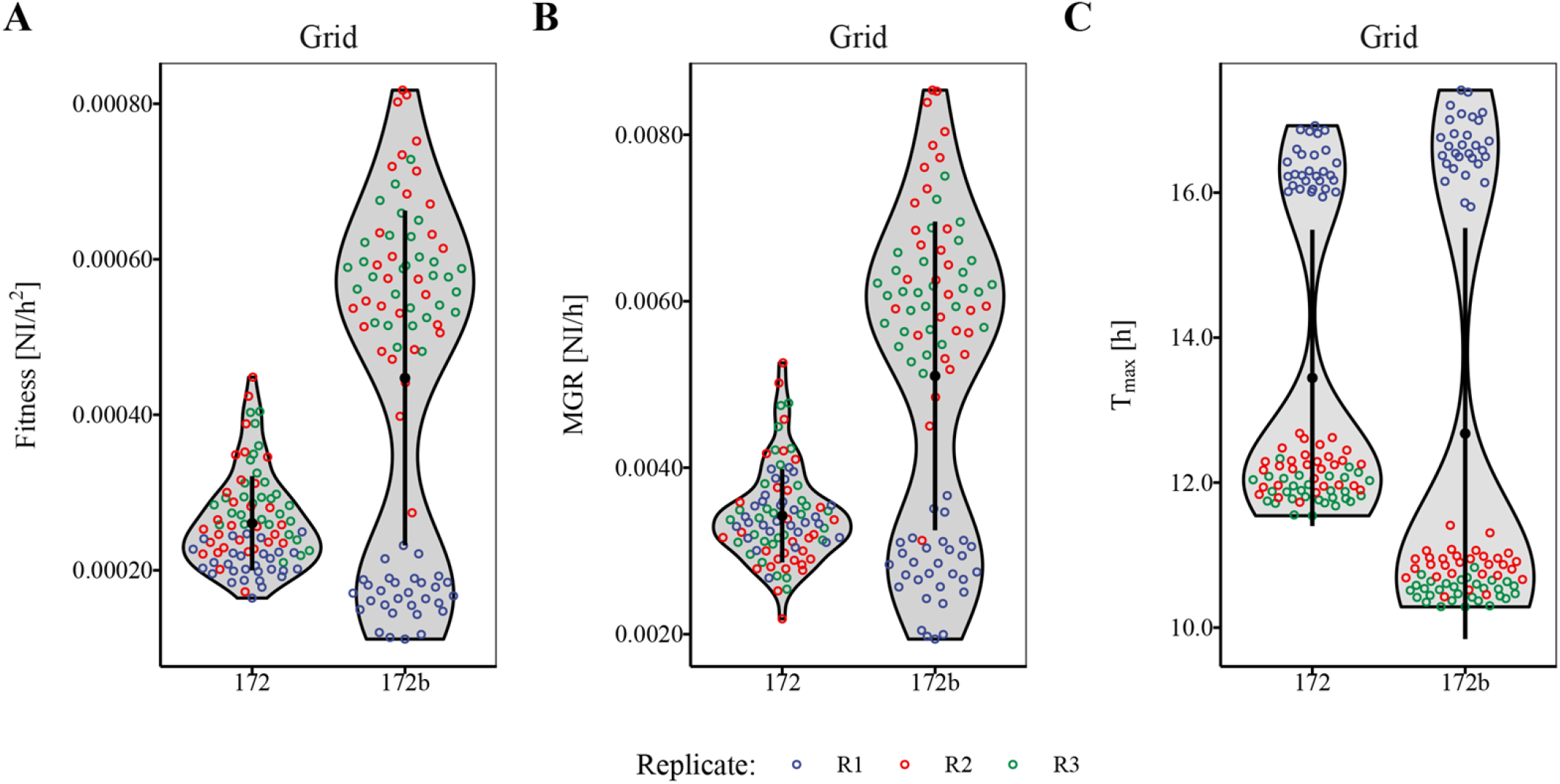
Fitness of vancomycin-sensitive *E. faecium* strain ST172 compared to the vancomycin-resistant (VRE) strain ST172b when grown on agar in a grid pattern.

## DISCUSSION

In this methods adaptation study, we adapted a previously established technique of quantitative fitness analysis (QFA) for use with important pathogenic bacterial strains with and without antimicrobial resistance. We were able to capture meaningful results from our baQFA adaptation, and detected differences in reproductive fitness between certain strains with and without antimicrobial resistance. We did not intend to answer the question whether the antimicrobial resistance alone was causal for the observed differences in fitness, and thus did not further examine isogenic strains with and without antimicrobial resistance mutations. Our newly developed BaColonyzer software showed a slightly larger scale range of normalised intensity (NI) values, illustrated by a linearly decreasing slope in the Bland-Altman plots. Although we did not intend for this specifically during development of the new algorithm, we considered the larger scale to potentially improve detection of smaller differences in fitness between strains.

Interestingly, we could observe a potentially important effect of interaction, when *S. aureus* strains JE2 and Cowan were grown competitively on one plate, with JE2 consistently reducing the fitness of Cowan while showing increased fitness during this competition. As at least in the early phase of growth, nutrient-competition effects seem to be minimal [12], the reason for this interaction could not yet be determined by baQFA. It could be hypothesised that with our baQFA method, bacterial interactions beyond competition for nutrients might be quantified, as they have been described similarly *in vivo* and *in vitro* [13,14]. Although we regard the results from our method adaptation work to only be of an exploratory nature, we could observe a substantially increased fitness in the methicillin resistant JE2 strain compared to the methicillin-susceptible Cowan strain, suggesting a different baseline-fitness of the underlying USA300 background in JE2 compared to Cowan, or the presence of compensatory mutations or other mechanisms of fitness gain. Interestingly, the *E. faecium* strain 172b, which differs from the 172 only by the presence of a vancomycin-resistance plasmid, was more fit in competitive baQFA, hinting at a possible fitness gain with the vancomycin-resistance plasmid or compensatory mutations.

Regarding cost, the baQFA specific materials can be obtained for less than $1,000, provided the lab is equipped with materials for bacterial growth (including an incubator, growth media, a replica plater and plates); if the consumer electronics are bought second hand, cost can be lowered to less than $500.

Our modified baQFA method has several strengths. First, it allows for an accurate determination of reproductive fitness with high throughput and at a very low cost, while being based on an established and validated technique. Second, the possibility of growing two strains in direct neighbourhood permits competitional effects to be investigated. Third, analysed bacteria are grown on semisolid medium, allowing for colony behaviour like quorum sensing as a possible factor in reproductive fitness. Finally, we have successfully used low-cost consumer electronics to automate the image capturing and data generation processes, making the method attractive for smaller labs, as well as research groups with a low budget for exploratory research, and investigators from low-resource settings.

Furthermore, our newly designed open-source software and R packages BaColonyzer and BaQFA are easy and fast to install. They also feature much faster analyses: BaColonyzer takes less than 1 second per image on an average laptop, about a fifth of the time the original Colonyzer software needed.

The adapted baQFA method also has several limitations. First, the BaColonyzer software, like its predecessor, is not yet able to correct for background colour changes during the time of growth, possibly making it difficult to accurately measure intensity for species like *Pseudomonas*, which produce coloured metabolites. Second, a direct measurement of bacterial species with a low optical signal (e.g., *Streptococcus spp.*), may not produce a high enough visibility to be adequately detected on photographic images, while the use of blood agar for enhanced contrast is not yet possible due to the change in colour over time. Third, we did not intend to compare different species and mainly focused on *Staphylococcus aureus* strains for data generation, thus deeming the resulting knowledge about the tested bacteria to be only of an exploratory nature. Finally, the absolute fitness measurements are difficult to universally standardize over different assays and laboratories. However, we expect the relative competitive fitness, derived from growing two strains on the same plate, to be less influenced by systematic factors, as they will affect both strains on the plate similarly, possibly making this relative measure more generalizable.

Overall, our low-cost and high-throughput baQFA method might be an important instrument for the prediction of epidemiologic persistence in the emergence of new resistant bacterial strains, as fitness has been proposed to be a major factor in the evolution of antimicrobial resistance [15].

For direct clinical application, a fast and inexpensive determination of fitness could be a useful co-factor to predict clinical outcomes in complex serious infections, or help clinicians in their choice and duration of antimicrobial therapies, adding reproductive fitness of the pathogen to the various host and pathogen factors being considered for optimal antimicrobial therapy concepts. Furthermore, baQFA might be used to evaluate the antibacterial activity of bacteriophages, as well as measuring phenomena like bacterial in-host evolution with loss of fitness after resistance acquisition and regain of fitness through compensatory mutations.

## CONCLUSION

In conclusion, we have adapted a quantitative fitness analysis method to be used with important pathogenic bacteria. We found that this adapted method provides valid fitness measures. The flexibility to automate the data generation with consumer electronics makes our method accessible for low-resource settings. This paves the way for further research focused on examining associations between reproductive fitness of bacterial pathogens and clinical outcomes, or epidemiological persistence of problematic bacterial strains, as reproductive fitness may be a major factor in these complex interactions [15].

## DECLARATIONS

### Ethics approval

As this study did not include any human participants, it was exempt from ethics approval according to the Swiss Human Research Act.

### Consent for publication

All authors have given their consent to the publication of this manuscript.

### Availability of data

The datasets used and analysed for our study are available from the corresponding authors upon reasonable request. All open-source software is available on GitHub (https://github.com/baQFA).

### Authors’ contributions

**Pascal M. Frey:** principal investigator, acquisition of funding, planning of the study, data collection, statistical analyses, drafting of the manuscript.

**Julian Baer:** data collection, statistical analyses, rewriting of BaQFA, drafting of the manuscript.

**Judith Bergadà Pijuan:** data collection, statistical analyses, BaColonyzer development, drafting of the manuscript.

**Conor Lawless:** development of the original QFA method, drafting and intellectual review of the manuscript.

**Philipp Bühler:** planning of the study, acquisition of funding, intellectual review of the manuscript.

**Roger D. Kouyos:** planning of the growth modelling, intellectual review of the manuscript.

**Kathrine P. Lemon:** planning of the study, acquisition of funding, intellectual review of the manuscript.

**Annelies S. Zinkernagel:** planning of the study, acquisition of funding, intellectual review of the manuscript.

**Silvio D. Brugger:** principal investigator, acquisition of funding, planning of the study, data collection, drafting of the manuscript.

## Acknowledgement

We would like to thank Dr. Federica Andreoni and Dr. Markus Hilty for their support of the study and critical review of this manuscript.

## FIGURE LEGENDS

Figure 1

After discarding the culture spots at the plate borders, the intensity measures for each time point are derived from time-lapse images using the custom build BaColonyzer software. Using the BaQFA package for R, growth curves and fitness measures can then be calculated from the normalised intensity measures. The blue dotted vertical line marks the point of maximum growth rate.

Figure 2

The conceptual design of the BaColonyzer algorithm is described in more detail in Supplement A.

Figure 3

The relationship of BaColonyzer normalised intensity (NI) measurements and colony forming units per milliliter (CFU/ml) counts was approximately log-linear for each strain, and showed some variation between the different strains. The linear regression found a strong association of NI and CFU counts (p < 0.01).

Figure 4

*S. aureus* strains JE2 and Cowan grown on agar either alone (Single) or together in a grid where each JE2 has only Cowan as direct neighbours during growth. Fitness (Panel A) is expressed as the maximum growth rate (Panel B) divided by the time to maximum growth (Panel C). Abbreviation: NI = Normalised Intensity.

Figure 5

Similar to *S. aureus*, baQFA showed that antibiotic resistance in *E. faecalis* ST172b was associated with a gain in fitness, compared to the closely related vancomycin-susceptible ST172. Fitness (Panel A) is expressed as the maximum growth rate (Panel B) divided by the time to maximum growth (Panel C). Abbreviation: NI = Normalised Intensity.

## Supplement A: Image Segmentation Algorithm

### Grid Location

BaColonyzer starts the analysis by finding the location of the grid. For that, it imports the grayscale version of the last image in the time-lapse series, which is used to generate a histogram of colours. Colours are represented as intensity values ranging from 0 to 255.

Since the last image contains well-defined colony spots, the histogram usually shows two high peaks: the highest peak corresponds to the colour of the agar while the second peak represents the colour of the colonies. As a quality check, BaColonyzer ensures that the colour intensity of the colonies falls between 70% of the maximum colour of the picture (upper bound) and twice the colour of the agar (lower bound). Otherwise, it sets the colour intensity of the colonies to be within this range.

The intensity values of agar and spots are used to create an artificial image of the plate (template), for which users need to input the number of rows and columns of the grid. Resizing and changing the position of this template allows to find the best match within the actual grayscale image. This is achieved by computing the lowest normalised squared difference.

Once the best match is found, BaColonyzer is able to predict the location of the whole plate and to trim those parts of the image that might introduce noise (e.g. the borders). Image trimming, followed by automatic thresholding using Otsu’s binarization, allows to get the position of the spots and the agar.

### Image Analysis

Each of the images in series is imported in grayscale and trimmed based on the grid location that has been previously derived.

By default, colour intensities of the image are also normalised (divided by 255) to be in range 0-1. Alternatively, in order to compare different series of time-lapse images, BaColonyzer allows the users to provide a reference picture for the normalization. This must be an image showing a white and black area next to each other, and must be taken using the same camera settings as the image series. The reference image is imported in grayscale and the colour intensities are clipped at 1%-99% quantiles to exclude outliers or noise. The resulting minimum and maximum intensity values (black and white, respectively) are taken to calibrate the colour intensities of images using the expression below:

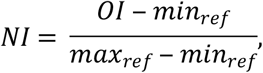

where NI are the normalised intensity values of each image, OI are the original values, min_ref_ is the intensity of the black colour, and max_ref_ is the intensity of the white colour.

Next, in order to ensure an accurate analysis of each colony, BaColonyzer divides the agar plate into smaller patches that only contain one colony spot. By default, the tool adjusts the intensity values of each patch by subtracting the mean intensity of the agar in this patch, thus correcting for colour differences within and between images.

At the end, BaColonyzer provides a result table with statistics of each colony, including colony mean, colony variance, background mean, and background variance. Results are stored as Output Data. Furthermore, in order to visually check that the grid location was achieved properly, BaColonyzer provides some output Images. These are binary images resulting from an Otus’s Binarization, which is performed after the trimming step to let the users check whether the grid location was successful.

## Supplement B

Bland-Altman plot comparing the new BaColonyzer software with the previous Colonyzer software for yeast. Each dot represents a single culture spot that was processed with both Colonyzer and BaColonyzer. The difference of those two measurements is represented by the y axis, while the x axis shows the mean of the measurements. With increasing mean intensity values, the old Colonyzer software systematically and linearly reported lower measurements than BaColonyzer.

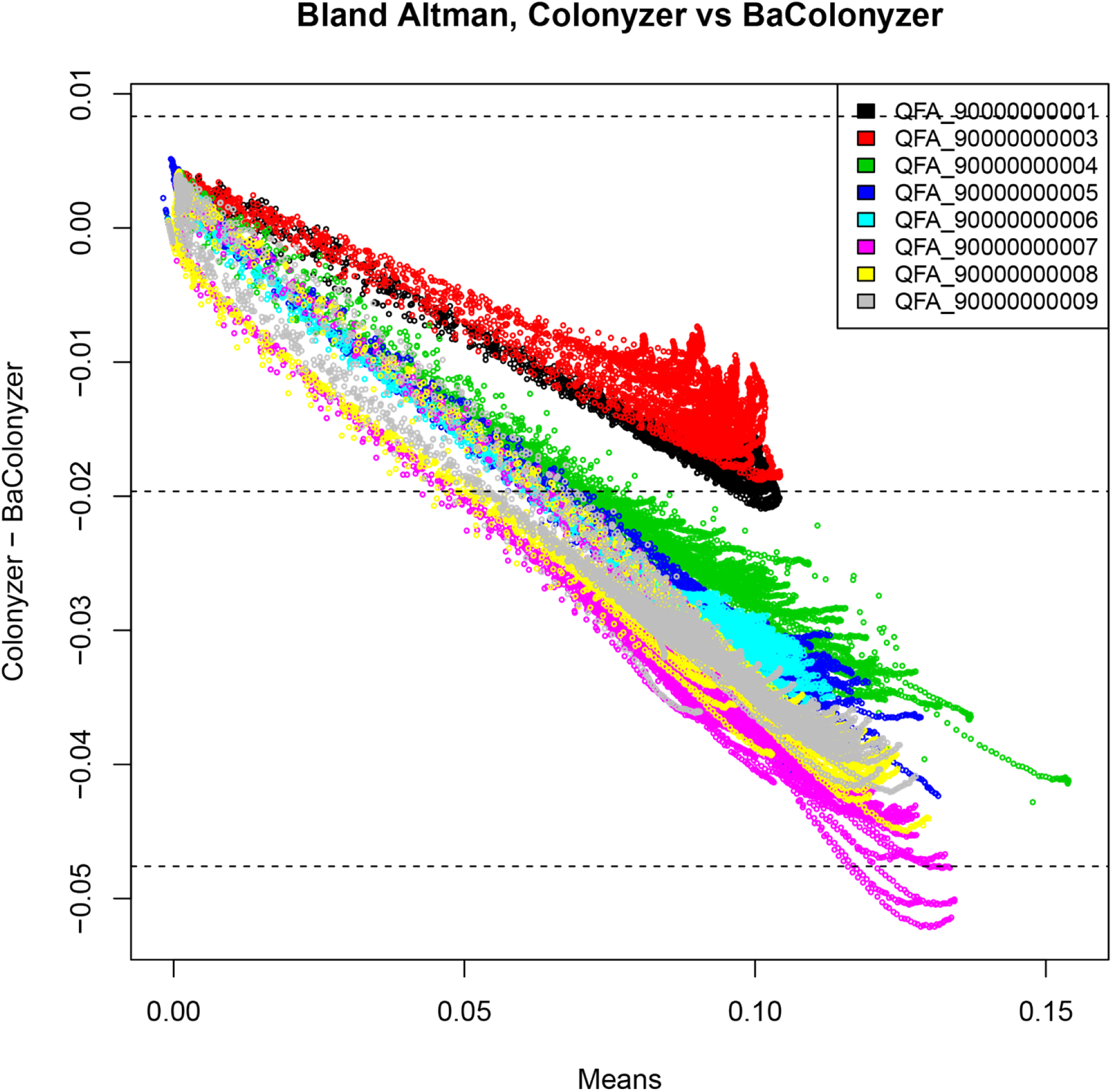

